# Arp2/3-mediated turnover of large clathrin lattices is regulated through the tyrosine kinase ACK

**DOI:** 10.64898/2026.06.12.731920

**Authors:** Mark A. Hazelbaker, Dennis J. Michalak, Mitchell T. Butler, Jordan R. Beach, Justin W. Taraska, James E. Bear

## Abstract

Clathrin mediated endocytosis (CME) is a vital cellular process that mediates cell signaling by controlling the internalization of extracellular cargo and activated receptors. Arp2/3-branched actin provides force to assist in membrane invagination and scission in CME. Loss of Arp2/3-branched actin in conditional *Arpc2* KO fibroblasts results in an increase of large arrested clathrin lattices (LACLs) visualized by live-cell TIRF imaging. Additional structural details of LACLs were revealed using electron microscopy and include an increase in arrested clathrin lattices of various curvatures. Numerous CME proteins have heightened levels of tyrosine phosphorylation at these LACLs in *Arpc2* KO cells. We identified the non-receptor tyrosine kinase ACK (Activated Cdc42-Associated Kinase) as a key upstream regulator of LACL turnover. CRISPR KO of ACK abrogates LACL tyrosine phosphorylation and impairs cells’ capacity to resolve LACLs. Our results support a model where ACK recruitment and activation at LACLs drives branched actin formation to help resolve large accumulations of arrested CME structures.

## INTRODUCTION

Branched actin networks play a central force-generating role in driving membrane remodeling and dynamic changes in cell shape which are critical for processes such as cell migration, endomembrane trafficking, and endocytosis (Campellone and Welch, 2010). The key element in this dynamic branched actin architecture is the Arp2/3 complex, a seven-protein complex that binds to existing actin filaments and nucleates a new daughter filament at ∼70° angle (Pollard, 2007). Arp2/3 branch formation is stimulated by nucleation promoting factors (NPFs) such as WAVE/SCAR, WASH, and N-WASP. NPFs integrate upstream signals from extracellular cues sensed by integrin adhesions and growth factor receptors, often through Rho-family GTPases, to spatially and temporally regulate the formation of branched actin (Rotty et al., 2013). However, much less is known about how Arp2/3-branched actin influences signaling networks in cells, particularly through its role in membrane remodeling. One such example is where loss of Arp2/3 complex triggers the upregulation of the senescence activated secretory phenotype (SASP) transcriptional signature via activation of nuclear factor κB (NFκB) (Wu et al., 2013).

Clathrin mediated endocytosis (CME) is a highly conserved process by which cells internalize plasma membrane receptors and associated cargo (Watts and Marsh, 1992; Bitsikas et al., 2014). Prototypical CME involves invagination of the membrane through a combination of a clathrin triskelion lattice, adaptors, dynamin and Arp2/3-branched actin (Kaksonen and Roux, 2018). These proteins are recruited to the membrane in groups with tightly coordinated temporal precision (Taylor et al., 2011). While the role of many of these proteins in CME is fairly well understood, some molecular players such as ACK (activated CDC42-associated kinase) remain understudied. In addition to canonical invaginating pits, cells can also form larger clathrin lattice structures that do not invaginate and are longer lived, often referred to as clathrin plaques (Saffarian et al., 2009; Lampe et al., 2016). When visualized using electron microscopy, these plaque-like structures display a heterogenous array of flat clathrin lattices (FCLs)(Heuser, 1980; Maupin and Pollard, 1983; Sochacki et al., 2021). FCLs have been suggested to function as signaling platforms by maintaining active receptors at the membrane for an extended times (Grove et al., 2014; Leyton-Puig et al., 2017; Baschieri et al., 2018; Alfonzo-Méndez et al., 2022). In myocytes and osteoblasts, FCLs are found at costameres and podosomes, respectively (Akisaka et al., 2008; Vassilopoulos et al., 2014). At these structures, FCLs may play a structural role, providing the providing a link between the cytoskeleton and the membrane. Loss of these FCLs in differentiated myotubes was shown to impair muscle integrity by disrupting sarcomere attachments to costameres (Vassilopoulos et al., 2014).

The factors that regulate the formation and dynamics of FCLs are not well understood. Alternative splicing of the clathrin heavy chain transcript to include exon 31 leads to increased formation of FCLs (Moulay et al., 2020). Other data indicate that integrins can regulate FCLs. For example, β1 integrin signaling negatively regulates FCL accumulation (Hakanpää et al., 2023). Conversely, FCLs colocalize with β5 integrin, forming reticular adhesions during mitosis (Lock et al., 2018), and stimulation of serum-starved cells with EGF produces an increase in FCLs that is dependent on EGFR, Src kinase, and β5 integrin (Alfonzo-Méndez et al., 2022). However, this effect is transient, with the levels of FCLs decreasing to baseline after one hour. The turnover of FCLs is dependent on the Arp2/3 complex as shown by the pharmacological inhibition of Arp2/3 inducing the formation of FCLs (Yang et al., 2022). Upon washout of the Arp2/3 inhibitor, branched actin forms at the perimeter of FCLs, specifically at breaks or edges of the lattice, where it is proposed to provide a force necessary for the completion of the process of clathrin lattice reshaping into a clathrin-coated vesicle. However, the mechanism by which Arp2/3 activity is regulated at non-typical clathrin lattices is unknown. Through an unbiased proteomic approach, we have discovered an ACK-dependent signaling pathway, upstream of Arp2/3, which is critical for FCL regulation.

## RESULTS AND DISCUSSION

### Loss of Arp2/3-branched actin causes accumulation of aberrant clathrin structures

To test how the loss of the Arp2/3-branched actin networks perturbs cellular regulatory processes, we used an unbiased proteomic approach. For these studies, we used a previously developed conditional *Arpc2* KO mouse dermal fibroblast cell line- JR20 cells (Rotty et al., 2017). This cell line has a tamoxifen inducible Cre recombinase (CreERT2) that, upon activation, excises exon 8 of the *Arpc2* gene, creating a mis-sense mutation that leads to loss of the ARPC2 (p34Arc) protein and all functional Arp2/3 complexes. To examine how signaling cascades change without Arp2/3, we subjected the *Arpc2* KO JR20 cells and matched WT controls to tyrosine phospho-proteomic analysis (**Fig. 1A**). Briefly, tyrosine phosphorylated peptides were enriched from trypsin digested cell lysates using a phospho-tyrosine (pTyr) antibody mix prior to analysis via liquid chromatography tandem mass spectrometry (LC-MS/MS). To our surprise, gene ontology analysis revealed that numerous proteins in the clathrin mediated endocytosis pathway, including the clathrin heavy chain and AP2, were hyper-tyrosine phosphorylated in *Arpc2* null cells (**Fig. 1B, Supplemental Table 1**). In addition, this list contains several proteins known to promote branched actin formation at sites of clathrin endocytosis, like the actin nucleation promoting factors (NPFs) N-WASP and WIP (Munn, 2001; Sun et al., 2017).

**Figure 1:**
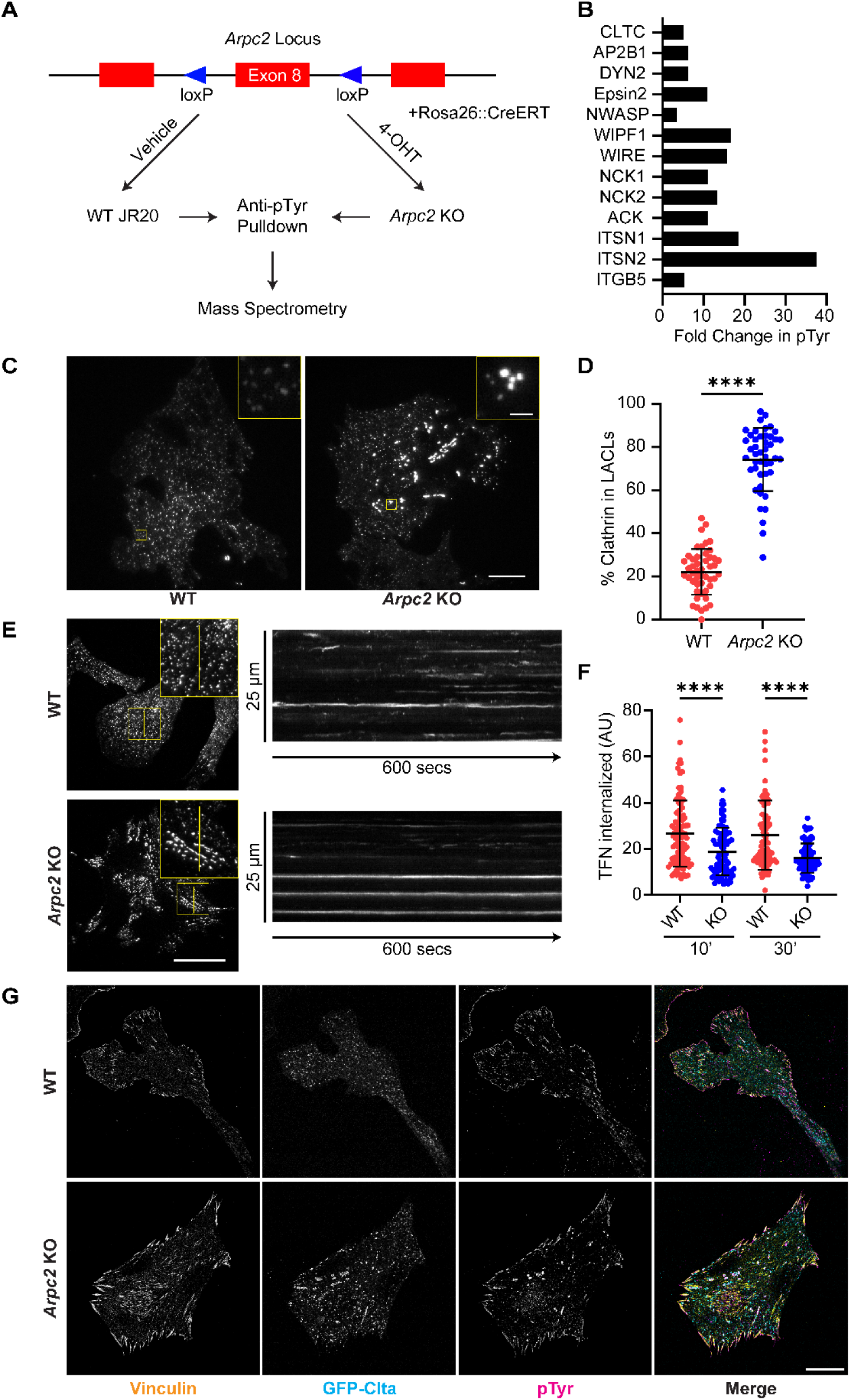
Loss of Arp2/3 complex leads to increased tyrosine phosphorylation of clathrin-mediated endocytosis pathway components. (A) Schematic representation of *Arpc2* KO phospho-tyrosine proteomic study workflow. (B) Average fold increase in tyrosine phosphorylation of CME related proteins across all tyrosine phosphorylation sites in *Arpc2* KO:WT cells. (C) Representative TIRF images of GFP-Clta in WT and *Arpc2* KO cells plated on 10 μg/mL FN for 48 hrs. Scale bar, 20 μm. (D) Percentage of GFP-Clta within LACLs vs all clathrin structures observed in TIRF field, by area, for parent (n=48) and *Arpc2* KO (n=45) from N=3 replicate experiments. (E) Example kymographs for WT and *Arpc2* KO cells. Scale bar, 20 μm. (F) Transferrin internalized by WT (n=97, 81) and *Arpc2* KO (n=79, 80) cells after 10 or 30 minutes from N=3 replicate experiments. (G) Immunofluorescence of vinculin, clathrin, and phosphotyrosine in WT and *Arpc2* KO cells. Scale bar, 20 μm. Welch’s t tests were performed for graphs D and F. ****p<0.0001. Error bars represent mean ± SD.

To determine how loss of Arp2/3 and the resulting changes in tyrosine phosphorylation affected clathrin dynamics, we imaged cells stably expressing GFP-tagged clathrin light chain (GFP-Clta). In WT cells, clathrin predominantly forms small, fairly uniform punctate structures when imaged via TIRF microscopy (**Fig.1C**), consistent with a combination of clathrin pits and small FCLs. In contrast, *Arpc2* null cells form many large clathrin structures that are heterogeneous in both size and shape in addition to the smaller structures similar to the WT cells. To ensure that this phenotype was not due to the expression of tagged clathrin, we also stained for endogenous clathrin heavy chains in cells without the exogenously tagged light chain and observed similar structures (**Supplemental Fig.1A**). In order to evaluate the dynamics of these clathrin structures, we imaged WT and *Arpc2* KO cells for 10 minutes. Kymographs of WT JR20 cells show that while a few clathrin structures are static, most exhibit dynamic behaviors often associated with CME events (Saffarian et al., 2009), appearing and disappearing during the imaging window (**Fig. 1E**). In contrast, the larger structures observed in *Arpc2* KO cells are highly stable over the 10 minutes. To compare the dynamics of the small, transient structures between WT and *Arpc2* KO cells, we employed an analysis pipeline to track diffraction-limited spots in our movies and measure their lifetimes (**Supplemental Fig. 1B**) (Aguet et al., 2013). Loss of Arp2/3 complex did not substantially affect the lifetime of CME events associated with the smaller, more transient structures we observed. To quantify the overall change in clathrin architecture, we segmented all clathrin structures in our images and saw a significant increase in the accumulation of large clathrin structures in the *Arpc2* null cells (**Fig. 1D**). We used a threshold of over 500 nm in diameter to classify these aberrant structures, which we will agnostically refer to as Large Arrested Clathrin Lattices (LACLs) for the remainder of this study.

To determine how the altered clathrin architecture affected functional endocytosis in the *Arpc2* null cells, we performed a transferrin uptake assay. We added fluorescently labeled transferrin to cells, fixed them at several timepoints, and measured the intensity of internalized transferrin across the whole cell. In comparison to WT cells, *Arpc2* null cells demonstrated an impaired ability to endocytose transferrin at both 10- and 30-minute timepoints (**Fig. 1E**). This observed decrease in uptake in the *Arpc2* null cells is consistent with previous studies (Almeida-Souza et al., 2018; Miller et al., 2018) and could reflect aborted uptake events, clathrin machinery being sequestered in LACLs, or a combination of both.

Next, we sought to determine the phospho-tyrosine content of the LACLs. Consistent with the proteomic data, staining with anti-pTyr shows that LACLs have high phospho-tyrosine levels (**Fig. 1F**). This staining is comparable to pTyr levels observed at focal adhesions, while anti-pTyr staining of the small clathrin puncta in either the *Arpc2* null or WT control cells was negligible. Therefore, we concluded that the increase of tyrosine phosphorylation on multiple CME proteins was occurring predominantly at LACLs and was not increasing generally at all clathrin structures within the cell. From this, we hypothesized that the hyper-tyrosine phosphorylated CME proteins were accumulating at the aberrant LACLs in Arp2/3 null cells in an attempt to resolve these frustrated structures by activating the absent Arp2/3.

To characterize clathrin morphology at nanometer-scale resolution in the absence of Arp2/3, we used platinum replica electron microscopy (PREM) on unroofed WT and *Arpc2* null cells (**Fig. 2A and B**). This approach allowed us to determine the structure and distribution of individual clathrin structures, which appear as distinct honeycomb patterned lattices, at the plasma membrane. Various levels of lattice curvature were observed and defined for quantification into three categories as 1) flat (little to no observable curvature), 2) domed (curved lattice with visible edges), and 3) spherical (curved with no visible lattice edge). Cortical actin networks were seen across unroofed membranes, appearing as 15 nm filaments that ranged from dense bundles to finer strands that sometimes branched. Consistent with loss of Arp2/3 activity, *Arpc2* null cells appeared to contain fewer putative branched filaments than WT controls, although this is difficult to assess definitively from 2D PREM projections.

**Figure 2:**
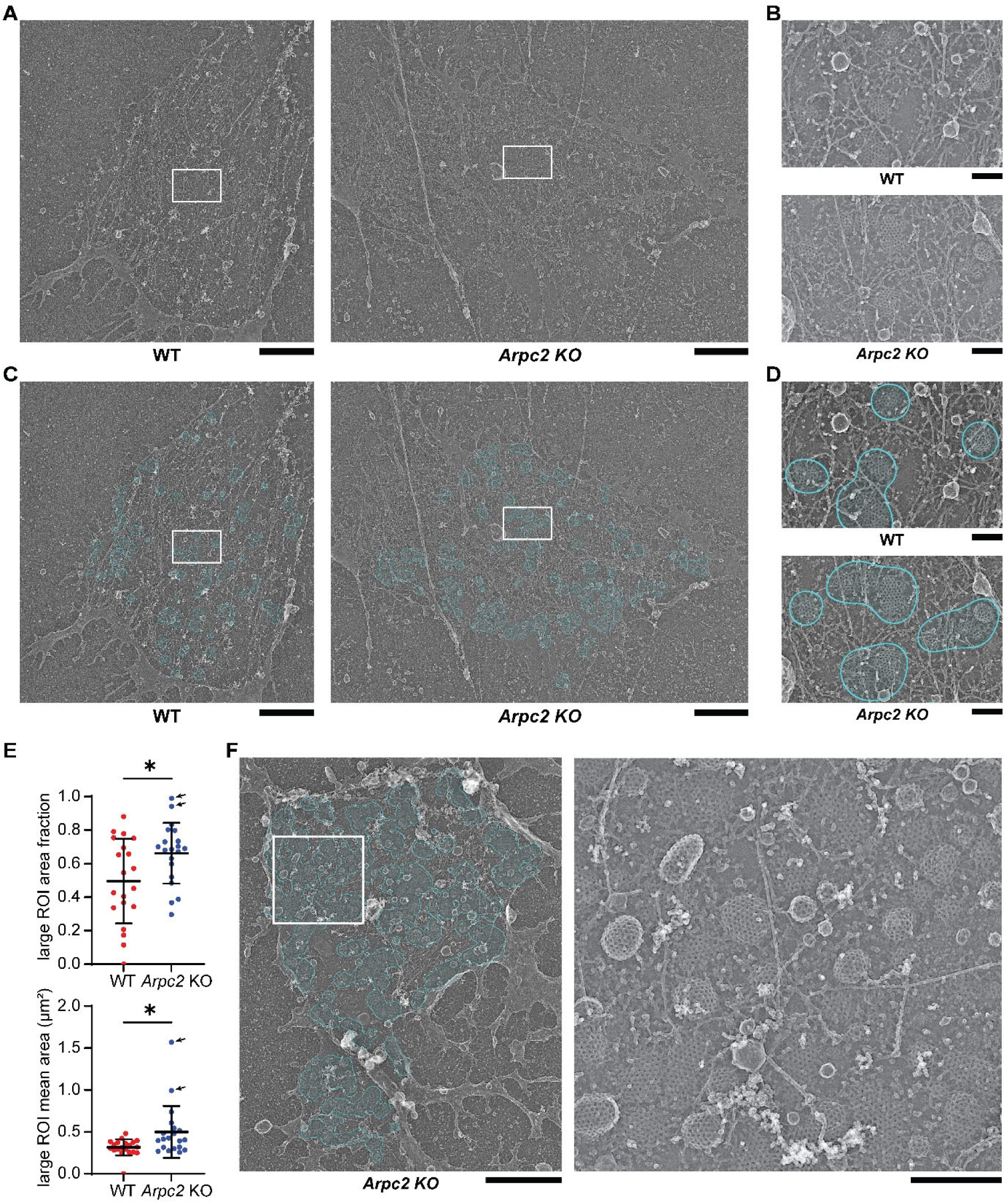
Loss of branched actin increases the size and proportion of large clathrin-enriched regions observed by PREM. **(A)** Representative PREM montages of isolated membranes generated by unroofing WT JR20 and *Arpc2* KO JR20. White boxes indicate regions shown in B. Scale bar, 2 µm. **(B)** Magnified views of the boxed regions in A. Scale bar, 250 nm. **(C)** Clathrin ROIs, cyan, derived from Gaussian blurred segmentations. White boxes indicate regions shown in D. Scale bar, 2 µm. **(D)** Magnified views of the boxed regions in C. Scale bar, 250 nm. **(E)** Quantification of clathrin ROIs from WT and *Arpc2* KO JR20 montages. Large ROI area fraction was calculated as the area occupied by ROIs with equivalent diameter >= 500 nm divided by the total clathrin ROI area. Mean large ROI area is shown in µm². Each point represents one montage; bars indicate mean ± SD. Arrows indicate *Arpc2* KO montages displaying the extreme phenotype. **(F)** A representative *Arpc2* KO membrane, left, with the extreme phenotype. Clathrin ROIs are outlined and shaded in cyan. The white box indicates the region shown in the right image. Scale bars, left and right, 2 µm and 500 nm, respectively.

To compare differences in clathrin at WT and *Arpc2* null cell membranes, we first segmented clathrin structures, classifying each as either flat, domed, or spherical. To more easily compare with TIRF-scale fluorescence and account for any error in finding the edge of the clathrin lattice, segmentations were pooled into a single class and Gaussian-blurred (σ = 90 nm). After applying a threshold, the resulting ROIs were used for quantification of clathrin abundance (**Fig. 2C and D**; **Supplemental Fig. 2A**). Similar to the fluorescence data, large ROIs were defined as having an equivalent diameter of ≥500 nm. Using this workflow, *Arpc2* null montages showed a higher fraction of clathrin ROI area contained within large ROIs and a larger mean area of these large ROIs compared with WT controls (**Fig. 2E**). These effects were robust across a range of parameters, indicating that the increased large-ROI clathrin area fraction was not specific to a single image-processing methodology (**Supplemental Fig. 2B**).

To determine if *Arpc2* disruption affected the process of clathrin curvature, we next compared two segmentation classes: flat and curved (where curved are objects pooled from domed and spherical classes) (**Supplemental Fig. 2C and D**). Of note, our method of segmenting clathrin into distinct units based on curvature results in a lattice with both curved and flat subregions being treated as more than one object. This analysis showed increased clathrin coverage in *Arpc2* null cells was driven primarily by an increase in the density of curved structural units. Interestingly, the average size of individual clathrin structural units remained constant across treatments. Thus, loss of Arp2/3 does not appear to increase the size of an individual flat or domed structural unit of clathrin.

Throughout our imaging datasets, a subset of *Arpc2* null cells displayed an extreme phenotype not observed in WT controls (**Fig. 2F**). In such cells, numerous flat and curved patches of clathrin are organized across the membrane as a network of extended clumps. These correspond to the upper range of the large-ROI quantified in **Fig. 2E** and contained densely packed, sometimes interconnected flat and curved clathrin structures spanning broad regions on the plasma membrane. These data indicate that Arp2/3 disruption stalls clathrin assembly at a stage following inward curvature but before separation of these vesicle-forming units from the plasma membrane (Sochacki et al., 2021; Lampe et al., 2016).

### Identifying factors involved in the tyrosine phosphorylation of clathrin related proteins

Previous studies showed that ß1-containing integrins (ITGB1) are critical to prevent the formation of excessive FCLs (Hakanpää et al., 2023). To determine whether high tyrosine phosphorylation is an inherent property of all large clathrin lattices, we used immunofluorescence to examine clathrin structures in an *ITGB1* null cell line (GD25) (Fässler et al., 1995). As expected, GD25 cells have a large proportion of their clathrin in LACLs vs. smaller structures (**Fig. 3A and C**). Interestingly, these LACLs did not co-stain with phospho-tyrosine, indicating that all LACLs do not inherently display high tyrosine phosphorylation levels. To confirm that this phenotype was a consequence of lack of ITGB1 signaling, we rescued these cells by stably expressing exogenous ITGB1-GFP. Expression of ITGB1A, the ubiquitously expressed ITGB1 isoform, reduced LACL levels (**Fig. 3B and C**). Interestingly, exogenous expression of GFP tagged ITGB1D, an isoform which is expressed in striated muscle and localizes to costameres, failed to decrease LACL levels. From this, we conclude that ITGB1A, and specifically its C-terminal tail, regulates LACL stability. Using this system, we confirmed our results with the *Arpc2* KO fibroblasts by treating ITGB1A-GFP expressing GD25 cells with CK666, an Arp2/3 small molecule inhibitor. Inhibition of branched actin in these cells resulted in an accumulation of LACLs containing notable phospho-tyrosine labeling (**Fig. 3D and E**). Together, these data support the idea that the tyrosine phosphorylation at LACLs is a signal for LACL turnover, downstream of ITGB1, and only accumulates if they cannot resolve, such as in the absence of Arp2/3 activity.

**Figure 3:**
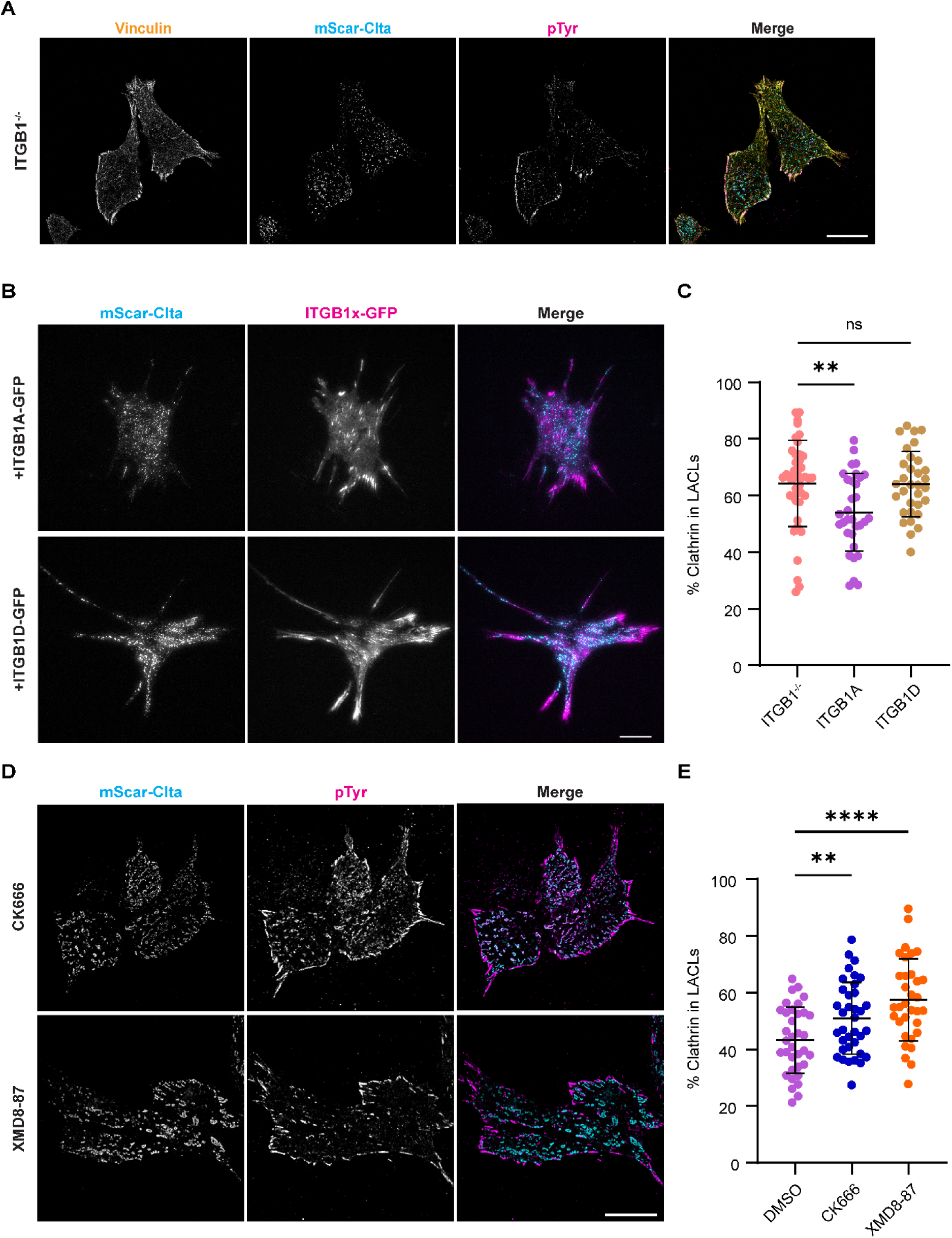
ITGB1 negatively regulates clathrin accumulation by promoting phosphotyrosine signaling at LACLs. **(A)** Immunofluorescence of vinculin, clathrin, and phosphotyrosine in ITGB1^-/-^ (GD25) cells. Scale bar, 20 μm. **(B)** Representative TIRF images of GD25 cells expressing mScar-Clta and either GFP-ITGB1A or GFP-ITGB1D. Scale bar, 20 μm. **(C)** Percentage of mScar-Clta within LACLs vs all clathrin structures observed in TIRF field, by area, for ITGB1 null (n=42), ITGB1A (n=32), and ITGB1D (n=32) expressing cells from N=3 replicate experiments. **(D)** Immunofluorescence of clathrin and phosphotyrosine in ITGB1A rescued cells treated with CK666 or XMD8-87 for 40 hours. Scale bar, 20 μm. **(E)** Percentage of mScar-Clta within LACLs vs all clathrin structures observed in TIRF field, by area, for ITGB1A cells treated with DMSO (n=35), CK666 (n=36), or XMD8-87 (n=33) from N=3 replicate experiments. Welch’s t tests were performed for graphs C and E. **p< 0.005, ****p<0.0001. Error bars represent mean ± SD.

While our findings using GD25 cells highlight an important link between ITGB1 and Arp2/3-mediated LACL turnover, we wanted to investigate and verify a similar mechanism is in place in our JR20 fibroblasts. Upon plating *Arpc2* null cells on either 10 µg/ml fibronectin (FN)-coated glass or 2% BSA-coated glass, we observed that cells plated on BSA coated glass had very little anti-pTyr signal relative to cells plated on FN, consistent with the idea that ITGB1 signaling is a strong upstream signaling input (**Supplemental Fig. 3**). To determine if serum affected tyrosine phosphorylation of LACLs, we incubated cells in low (0.2% FBS) or full (10% FBS) serum conditions after they had adhered. Cells in low serum had diminished levels of pTyr signal on FN-coated glass compared to full serum, and the pTyr signal in low serum was negligible when plated on BSA-coated glass. These data indicate that tyrosine phosphorylation of LACLs has signaling inputs from both ECM via ITGB1 and factors contained in serum.

Having confirmed a role of ITGB1 signaling in LACL resolution, we next sought to identify the tyrosine kinase responsible for the observed phosphorylation of LACLs and cross-referenced our phospho-proteomic data with the existing clathrin signaling and dynamics literature. We identified Activated CDC42-associated Kinase (ACK), a non-receptor tyrosine kinase, as a possible candidate (Manser et al., 1993; Kan et al., 2023). This kinase is highly tyrosine phosphorylated in the *Arpc2* null cells (**Fig. 1B, Supplemental Table 1**), is known to interact with clathrin and other CME proteins, and is downstream of multiple cell surface receptors including integrins (Yang et al., 1999; Kato-Stankiewicz et al., 2001; Teo et al., 2001; Yokoyama et al., 2005; Galisteo et al., 2006). To evaluate ACK’s role in regulating tyrosine phosphorylation of LACLs, we used XMD8-87 to inhibit ACK in ITGB1A expressing GD25 cells. ACK inhibition resulted in increased numbers of LACLs, but importantly these LACLs lacked tyrosine phosphorylation (**Fig. 3F and G**). This suggests that ACK is an important regulator of both LACL formation and/or turnover, as wells as their tyrosine phosphorylation.

### The role of ACK in regulating clathrin dynamics

After identifying ACK as a putative regulator of LACLs, we expressed mScarlet-ACK in our JR20 system to observe how it localized relative to various clathrin structures. Live-cell TIRF imaging showed mScarlet-ACK^WT^ transiently interacting with smaller clathrin structures in WT JR20 cells, consistent with previous report (Taylor et al., 2011). In contrast, mScarlet-ACK^WT^ stably associated with LACLs in *Arpc2* null cells (**Fig. 4A**). To quantify this, we used intensity line scans of both LACLs and small clathrin puncta.

**Figure 4:**
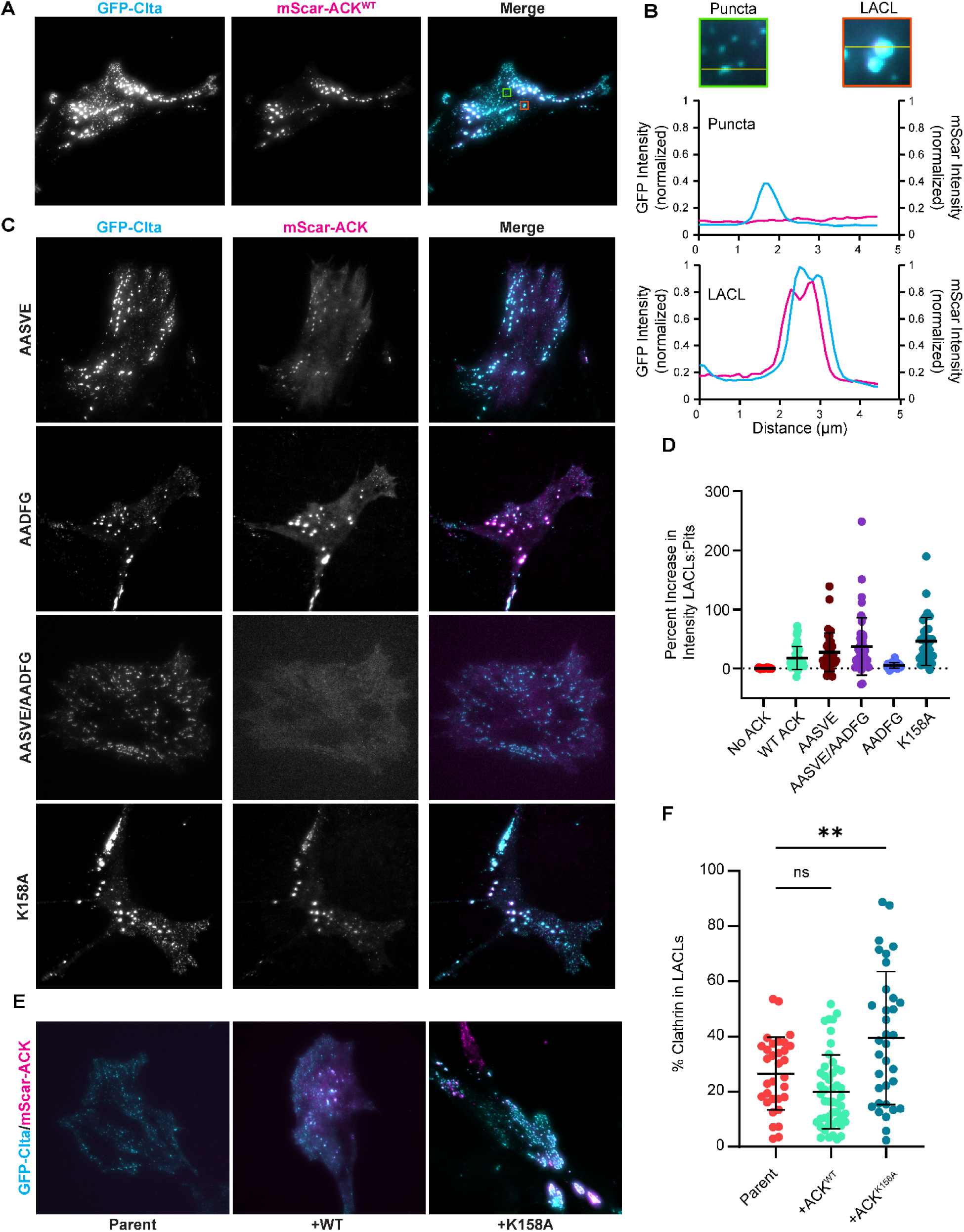
ACK preferentially localizes to LACLs. **(A)** *Arpc2* KO cells expressing GFP-Clta and mScarlet-ACK^WT^ were plated on BSA coated glass for 48 hours prior to imaging via TIRF microscopy. **(B)** Intensity profile of GFP and mScarlet across clathrin pits (blue inset) and LACLs (orange inset). **(C)** Representative TIRF images of *Arpc2* KO cells expressing GFP-Clta and mScarlet-ACK constructs, plated on BSA coated glass for 48 hours. **(D)** Percent increase in LACL mean mScarlet intensity compared to mean mScarlet intensity at pits for control (n=27), ACK^WT^ (n=38), ACK^AASVE^ (n=36), ACK^AADFG^ (n=45), ACK^AASVE/AADFG^ (n=27), and ACK^K158A^ (n=32) from N=4 replicate experiments. **(E)** Representative TIRF images of WT JR20 cells expressing GFP-Clta and mScarlet-ACK, plated on 10 μg/mL FN for 48 hours**. (F)** Percentage of GFP-Clta within LACLs vs all clathrin structures observed in TIRF field, by area, for parent (n=32), +ACK^WT^ (n=48), and +ACK^K158A^ (n=34) from N=3 replicate experiments. *p< 0.01. Error bars represent mean ± SD. Scale bars, 20 μm.

These show a robust co-localization between mScarlet-ACK^WT^ and GFP-Clta at LACLs, but very little at smaller clathrin structures (**Fig. 4B**). This selective accumulation of ACK at LACLs suggests that ACK is binding preferentially to LACLs. An alternative explanation is that after binding to LACLs, ACK is unable to dissociate from the stabilized lattices, despite being able to do so with clathrin pits (Taylor et al., 2011).

Next, we investigated how structural motifs within ACK determine its localization and function by creating a series of mutant versions of the kinase and expressing these in our JR20/Arpc2 KO system. We quantified the effects on localization by comparing the ratio of mScarlet-ACK intensity to GFP-Clta at LACLs vs pits. First, we mutated ACK’s two known clathrin binding motifs, ACK^496-500^ (LLSVE) and ACK^569-573^ (LIDFG) (Yang et al., 2001; Teo et al., 2001; Shen et al., 2011), by mutating the first two amino acids to alanine either separately (ACK^AASVE^ and ACK^AADFG^) or together (ACK^AASVE/AADFG^). In *Arpc2* null cells, the two individual clathrin binding mutants (ACK^AASVE^ and ACK^AADFG^) retained their preferential localization to LACLs (**Fig. 4C and D**). However, the loss of both clathrin binding motifs prevented association with all clathrin structures when visualized via TIRF imaging. From this, we concluded that while either clathrin binding motif is sufficient for proper localization, at least one is necessary, consistent with previous immunoprecipitation experiments (Shen et al., 2011). Next, we created a kinase dead construct by mutating the catalytic lysine at 158 to alanine (ACK^K158A^). In *Arpc2* null cells, this construct also retained localization at LACLs, indicating that kinase activity is not required for proper localization. Strikingly, when expressed in parental JR20 cells, mScalert-ACK^K158A^ induced accumulation of LACLs compared to control cells (**Fig. 4E and F**). This suggests that the kinase-dead ACK mutant can act as a dominant negative, possibly by outcompeting endogenous ACK and precluding tyrosine phosphorylation.

To more directly address the role of ACK in the regulation of clathrin dynamics, we used CRISPR to generate ACK knockouts (gene name *Tnk2*) in our JR20 cell line. From these, we plated two distinct KO clones alongside control cells (Cl.1 and Cl.6) on FN-coated glass and cultured in low serum conditions, which promotes detectible but tempered pTyr signaling at clathrin structures in *Arpc2* KO cells (Supplemental Fig. 3) and is more conducive to LACL formation than full serum conditions. Both clones showed a significant accumulation of LACLs compared to WT cells (**Fig. 5A and B**).

**Figure 5:**
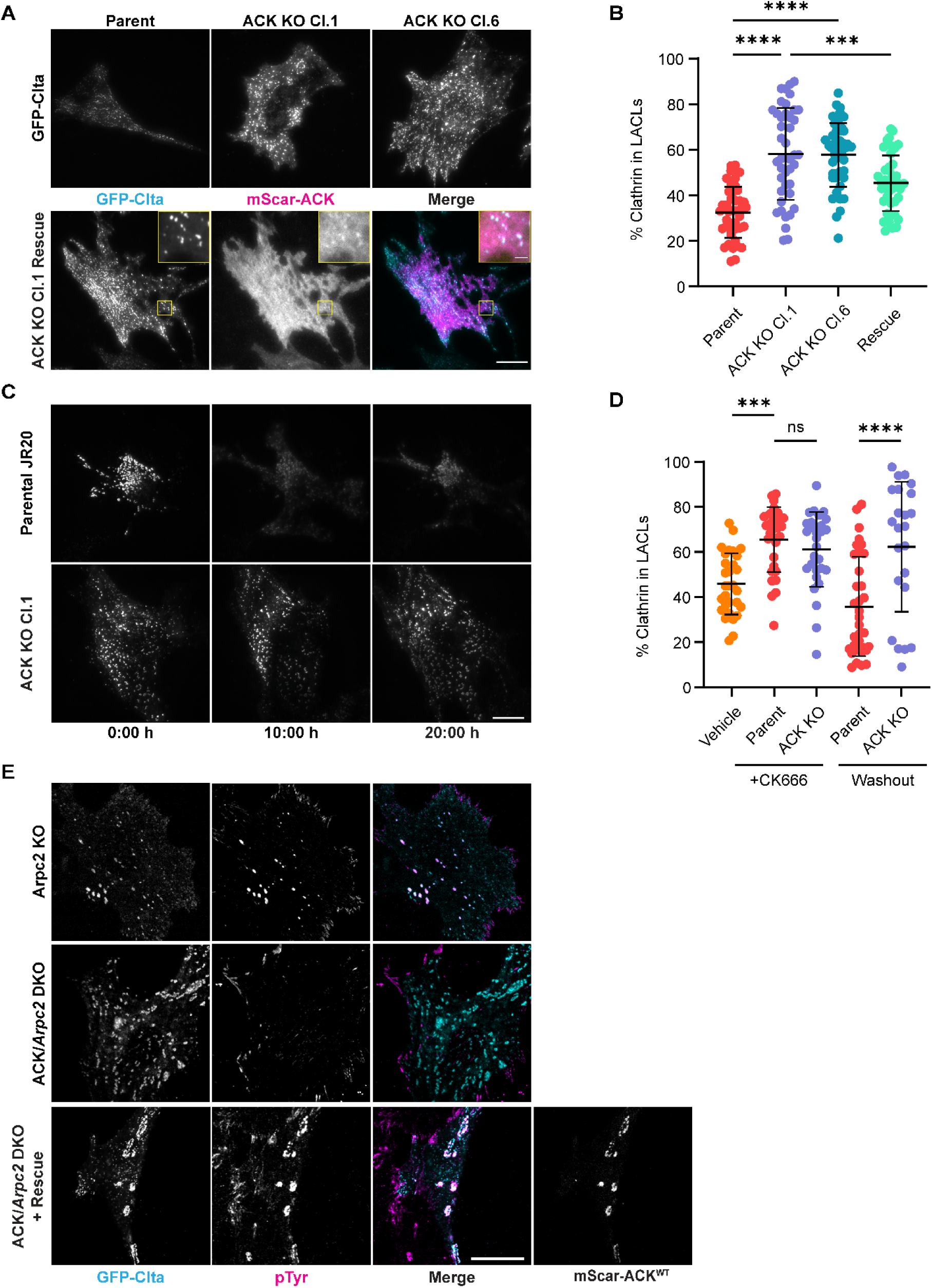
ACK is required for hyper tyrosine phosphorylation at LACLs and their efficient resolution. **(A)** Representative TIRF images of WT, ACK KO, and ACK^WT^ rescue cells 48 hours post plating. Scale bar, 20 μm. **(B)** Percentage of GFP-Clta within LACLs vs all clathrin structures observed in TIRF field, by area, for parent (n=43), ACK KO Clone1 (n=43), ACK KO Clone6 (n=48), and ACK KO Clone1 rescued with mScar-ACK^WT^ (n=43) from N=3 replicate experiments. **(C)** Montage of CK666 washout from WT and ACK KO Clone1 cells. **(D)** Percentage of GFP-Clta within LACLs vs all clathrin structures observed in TIRF field, by area, for WT cells treated with DMSO (n=30), WT (n=30) and ACK KO Clone1 (n=30) treated with CK666 for 40 hours (N=3 replicate experiments), and WT (n=34) and ACK KO Clone1 (n=24) cells 20 hours after CK666 washout. **(E)** Immunofluorescence of vinculin, clathrin, mScarlet-ACK, and phosphotyrosine in *Arpc2* KO, ACK/*Arpc2* DKO, and ACK/*Arpc2* DKO cells rescued with mScar-ACK^WT^. Welch’s t tests were performed for graphs B and D. ***p=0.0007, ****p<0.0001. Error bars represent mean ± SD.

This result corroborated our previous findings with both the pharmacological inhibition and the kinase-dead ACK, cumulatively serving as strong support for this kinase as a critical regulator of clathrin architecture. Furthermore, we are able to rescue this ACK KO phenotype with expression of mScarlet-ACK^WT^, which ameliorates LACL accumulation (**Fig. 5A and B**).

Since both the Arp2/3 complex and ACK are important for the regulation of clathrin dynamics, we sought to understand their functional interaction in this process. First, we used CK666 to inhibit Arp2/3 activity in WT and ACK KO cells and observed similar accumulation of LACLs in both cell types (**Fig. 5C-D**). Upon subsequent washout of CK666, WT cells quickly resolve most of their LACLs. However, in ACK KO cells, the LACLs are retained for up to 20 hours after drug wash-out (**Fig. 5D**). From this, we conclude that cells are capable of resolving LACLs after their formation and accumulation, and that this resolution is dependent upon both Arp2/3 and ACK. We hypothesized that ACK was responsible for most of the hyper-tyrosine phosphorylation observed in the *Arpc2* null cells. To test this, we plated *Arpc2* KO, *Arpc2*/ACK double KO, and *Arpc2*/ACK double KO cells rescued with mScarlet-ACK^WT^ on FN-coated glass in full serum conditions and stained for phospho-tyrosine (**Fig. 5E**). Based on our studies, this should be most optimal conditions for producing pTyr-enriched LACLs.

Indeed, in *Arpc2*KO cells, we saw robust pTyr of LACLs. However, in *Arpc2*/ACK double KO cells, LACLs formed but had no discernable anti-pTyr signal. Expression of mScarlet-ACK^WT^ in *Arpc2*/ACK DKO cells rescued pTyr labeling, which strongly co-localizes with the mScarlet-ACK signal at LACLs. Together, these results indicate that ACK is necessary for the tyrosine phosphorylation observed at LACLs and regulates LACL accumulation.

Previous work showed that Arp2/3-branched actin plays an important role in regulating clathrin mediated endocytosis (Moreau et al., 1997; Merrifield et al., 2004; Boulant et al., 2011; Sun et al., 2017; Kaksonen and Roux, 2018). In this study, we investigated the role of Arp2/3-branched actin in larger clathrin assemblies and the mechanism by which it is regulated. The presence of both flat and domed clathrin lattices in *Arpc2* KO cells (**Fig. 2A and B**) suggests that Arp2/3-branched actin is not strictly necessary to induce curvature of lattices. Supporting this view, we observed a greater number of curved clathrin units in *Arpc2* KO cells, while their average size was not detectably affected by Arp2/3 loss (**Supplemental Fig. 2D**). Rather, branched actin may be necessary to either prevent LACL formation or to facilitate their progression toward productive endocytosis. Our data show that, unlike WT cells, ACK KO cells fail to resolve CK666-induced LACLs upon washout of CK666. This suggests that Arp2/3-branched actin is involved in LACL resolution and that ACK specifically regulates Arp2/3’s role in LACL turnover after they have formed, consistent with the localization of ACK to LACLs.

Interestingly, ACK KO cells show a milder phenotype than *Arpc2* KO cells, suggesting that alternative pathways may also contribute to branched actin formation at CME sites and decrease LACL formation. One possible pathway includes intersectin-1/2 and CDC42, which has been shown to promote N-WASP activity during canonical CME (Hussain et al., 2001; McGavin et al., 2001).

An unresolved question arising from this work is how cells regulate LACL formation and turnover in cell type-specific physiological contexts such as in muscle cells. One intriguing potential mechanism for stabilizing LACLs in striated muscle is alternative splicing of CME related proteins, as is observed with CLTC and ITGB1 (Belkin et al., 1996; Moulay et al., 2020). These muscle-specific protein isoforms contain splicing-specific inserts that could affect the dynamics of the clathrin lattice itself or alter local signaling pathways inputs into the LACL resolution pathways in the case of ITGB1.

While our data confirmed previous reports that ITGB1A activity negatively regulate LACLs, the mechanism was unclear. Previously, it has been reported that ACK pY284 (an indicator of its activity) (Yokoyama and Miller, 2003; Kan et al., 2023) increased upon cell spreading in an ITGB1 dependent manner (Yang et al., 1999) and that this activation may require a Src family kinase (Linseman et al., 2001; Yokoyama and Miller, 2003; Chan et al., 2011). However, ACK kinase activity has not yet been shown to affect clathrin dynamics. Because ITGB1A is ubiquitously expressed, this proposed integrin-ACK pathway may be responsible for differentially modulating LACL levels in other cell types.

ACK has long been a promising candidate for targeted cancer treatment (Lawrence et al., 2015; Hodder et al., 2023) due to interactions with p53, Wwox, and p27, which lead to increased ubiquitination and subsequent degradation of these proteins (Mahajan et al., 2005; Xu et al., 2015; Peng et al., 2022), and its overexpression/activation in several types of cancer (Mahajan and Mahajan, 2013; Hodder et al., 2023). The ACK-mediated activation of the androgen receptor and AKT/PI3K pathway has made it a particularly high priority target in prostate cancer (Mahajan et al., 2007, 2010). Given our findings, it will be important to consider ACK’s role in regulating clathrin architecture and its potential effects on endocytosis in future targeting efforts.

## MATERIALS AND METHODS

### Cell culture

Cells were cultured in DMEM with 4.5 g/L D-glucose, L-glutamine and 110 mg/L sodium pyruvate (Gibco, Cat#: 11995-065) supplemented with 10% fetal bovine serum (FBS, Omega Scientific, Cat#: FB-01) at 37°C and 5% CO2. To passage, cells were washed with DPBS (Gibco, Cat#: 14190-144) and then detached using 0.25% trypsin-EDTA (Gibco, Cat#: 25200-072). To induce recombination at *Arpc2* locus to generate knockout cells, 2 µM 4-hydroxytamoxifen (Millipore Sigma, Cat#: H6278) was added to the low density JR20 culture media at least 7 days prior to an experiment.

For high-resolution TIRF and confocal imaging, glass bottom dishes (Cellvis, Cat#: D35-20-1.5N) were first coated with 10 µg/ml fibronectin (Corning, Cat#: 356008) or 2% bovine serum albumin (BSA, Fisher, Cat#: BP1600-100) in DPBS for 30 minutes at 37°C and washed 1x with DPBS prior to plating cells. For experiments that required low serum conditions, cells were initially plated in DMEM with 10% FBS for 4 hours, washed 1x with DPBS, and then incubated in DMEM with 0.2% FBS.

### Plasmid generation

To visualize clathrin in live cells, cells were transduced with mouse EGFP-Clta or mScarlet-Clta in the pLL5.0 lentiviral backbone (Vitriol et al., 2007) for stable expression. The Clta sequence was cloned from JR20 cDNA with SuperScript II Reverse Transcriptase (Invitrogen, #18064022) and inserted into a gateway entry vector (ThermoFisher K240020). A modified Gateway destination site and in-frame GFP sequence was moved from pCSdest-GFP (a kind gift from the Wallingford Lab) into lentiviral pLL5.0 plasmid using Gibson assembly, and the Clta CDS was introduced into this plasmid vector via Gateway LR reaction (ThermoFisher 11791020). The GFP sequence was exchanged with mScarlet via Gibson assembly. Intregrin Beta1-GFP pFB-Neo was a gift from Martin Humphries (Addgene # 69767) (Parsons et al., 2008). An integrin Beta1D gene fragment was synthesized (Twist Bioscience) and subcloned into pFB-Neo ITGB1A backbone with NEBuilder HiFi DNA Assembly Master Mix (NEB, #M5520A). Ack1-pmCherryC1 (Addgene plasmid # 27684, a gift from Christien Merrifield) (Taylor et al., 2011), was subcloned into the pLL5.0 mScarlet lentiviral backbone and the doxycycline inducible TLCV2 backbone (Addgene plasmid # 87360, a gift from Adam Karpf) (Barger et al., 2019). All PCR products were generated using Q5 High-Fidelity polymerase (NEB, #M0491) and assembled using NEBuilder HiFi DNA Assembly Master Mix (NEB, #E2621).

### ACK Mutagenesis Sequences

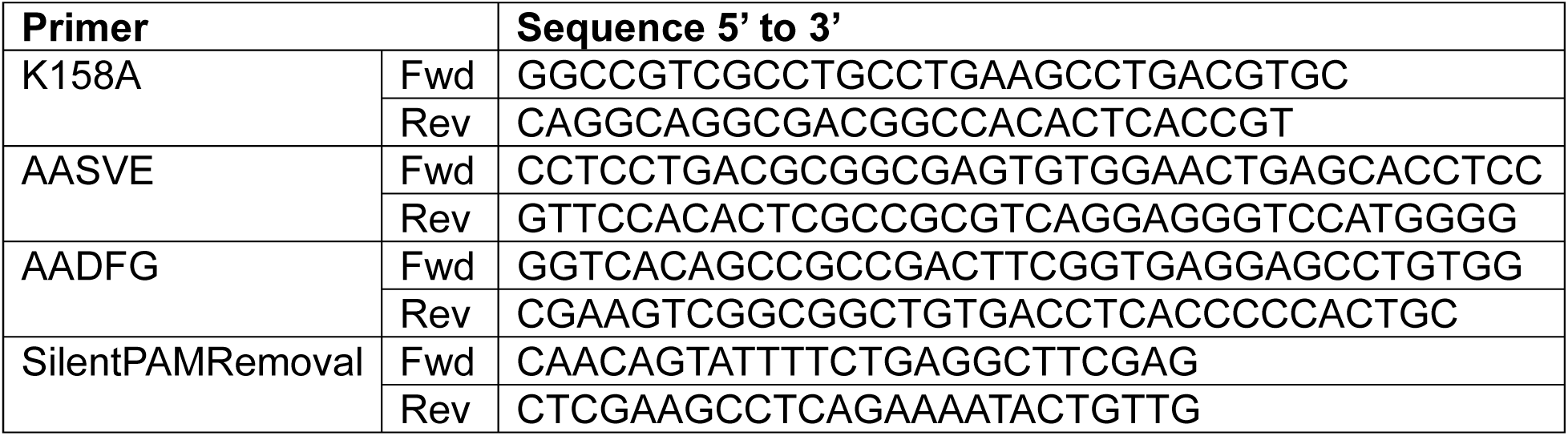

### Tyrosine phosphoproteomics

Trypsin digested WT and *Arpc2* KO cell peptides were immunoprecipitated using a pan-phosphotyrosine antibody (Cell Signaling, Cat#: 8803). Phosphotyrosine enriched peptides were then separated via liquid chromatography on a C18 reverse-phase PicoFrit capillary and analyzed via tandem mass spectrometry using an Orbitrap-Fusion Lumos (Thermo). Peptide spectra were identified using Comet (Eng et al., 2013).

Samples were run in duplicate (PTMScan, Cell Signaling). The fold change in tyrosine phosphorylation levels was averaged across all phosphotyrosine peptides detected for a given protein.

### Platinum replica electron microscopy (PREM)

Cells were grown on fibronectin-coated coverslips (Neuvitro, GG-25-1.5-fibronectin) for 24 hours in media at 37°C in 5% CO₂. Cells were briefly washed in unroofing buffer (30 mM HEPES, 70 mM KCl, 5 mM MgCl₂, 3 mM EGTA, pH 7.4) immediately prior to unroofing with an air pressure-driven syringe containing unroofing buffer supplemented with 2% paraformaldehyde. Unroofed cells were immediately transferred to DPBS containing 2% glutaraldehyde overnight at 4°C. Coverslips were then incubated in 0.1% tannic acid (in ddH₂O) for 20 minutes, washed with ddH₂O, and incubated in 0.1% uranyl acetate (in ddH₂O) for 20 minutes, with each step at 4°C. Coverslips were extensively washed with ddH₂O and dehydrated stepwise, as described previously (Sochacki et al., 2017; Sochacki and Taraska, 2017), at room temperature for 4 minutes per step. Coverslips were then critical point dried using an Autosamdri-815 (Tousimis) and transferred to a Leica EM ACE 900 for rotary shadowing with a platinum thickness target of 3 nm and a backing carbon layer target of 5.5 nm. Replicas were separated from coverslips using hydrofluoric acid and transferred to TEM grids for imaging on a JEOL 1400 with SerialEM.

Montages were stitched using the IMOD software suite (Mastronarde and Held, 2017). Each montage was either manually segmented as described previously (Sochacki et al., 2021) or automatically segmented with manual correction using PyCLEM (Sun et al., 2025). The edges of unroofed membranes were outlined, and clathrin structures were identified and classified into three morphological categories: flat (no visible curvature), domed (curved lattice with a visible edge), and sphere (curved with no visible lattice edge). For comparisons with quantification from TIRF data, segmentations were converted to a binary mask and Gaussian blurred with a blurring radius of 90 nm. These blurred masks were thresholded at 0.2, and ROIs were obtained from connected components. Large ROIs were defined as having equivalent diameters greater than 500 nm, in line with the TIRF analysis.

### Immunofluorescence and antibodies

Cells were fixed for 10 minutes using 4% paraformaldehyde (Electron Microscopy Sciences, Cat#: 15713) in DPBS at room temperature. Cells were permeabilized using 0.2% Triton-X (ThermoFisher, #BP151) in DPBS for 5 minutes and blocked in 5% BSA in PBS for 30 minutes at room temperature. For staining endogenous clathrin heavy chain, cells were instead permeabilized with 0.2% saponin (Sigma, #S4521-10G) in DPBS for 5 minutes. Primary and secondary antibodies were diluted in 5% BSA and incubated for 1 hour at room temperature. Cells were washed 3x in DPBS after each step.

Antibodies were obtained from Santa Cruz (Clathrin Heavy Chain [12734]), Cell Signaling (Phospho-Tyrosine [8954S]), Sigma-Aldrich (hVin1 [V9131]), Jackson ImmunoResearch (Cy5 Goat anti-Rabbit [111-175-148] and FITC Donkey anti-Mouse [715-095-150]) and ThermoFisher (DyLight 405 Goat anti-Mouse [35500BID]).

### Microscopy and image analysis

TIRF images were acquired using Olympus 60x Apo N (1.49 NA) or Olympus 60x UPlanApo (1.50 NA) TIRF objectives with an Orca-Flash4.0 (Hamamatsu) camera on an Olympus IX81 or IX83 microscope. Confocal images were acquired using a 63x

PlanApo (1.4 NA) objective with a Zeiss LSM800. Widefield fluorescence images were captured on a Zeiss LSM800 with a 20x PlanApo objective and an Orca-Flash4.0LT (Hamamatsu) camera. For all live cell imaging, cells were incubated at 37°C and 5% CO2 using a Tokai Hit stage top incubator.

#### LACL accumulation analysis

LACL accumulation was quantified using a CellProfiler pipeline. First, individual cells were segmented using Otsu’s method with a global threshold after applying a gaussian filter across the TIRF image. Segmented cells were then used to mask the original image. Clathrin structures within cells were then segmented using Otsu’s method with an adaptive window of 50 pixels. Structures with a diameter above 500 nm were classified as LACLs. The percentage of clathrin in LACLs was calculated by dividing the integrated area of LACLs by the integrated area of all segmented clathrin structures in the TIRF field (LACLs and pits/FCLs with a diameter below 500 nm).

#### Clathrin lifetime analysis

Two days before imaging, cells were plated on 10 µg/ml FN coated glass in DMEM with 10% FBS. WT and *Arpc2* KO cells were imaged via TIRF every second for 10 minutes, and the Multiple Kymograph plugin in ImageJ was used to create kymographs along a 25 µm line from the movies. Lifetimes of endocytic events were quantified using the MATLAB package cmeAnalysis, which detects and tracks diffraction-limited spots (Aguet et al., 2013).

#### ACK localization analysis

*Arpc2* KO cells stably expressing EGFP-Clta and mScar-ACK^WT^ were plated on 10 µg/ml FN coated glass for 48 hours prior to imaging via TIRF microscopy. Intensity profiles for both channels were generated along a linear region of interest using the Plot Profile plugin in ImageJ. To quantify ACK accumulation on LACLs, the average mScarlet intensity of individual clathrin structures was measured. The percent increase in average mScarlet fluorescence of both LACLs compared to smaller structures was calculated per cell.

### CRISPR Knockout

To generate CRISPR knockouts of ACK, the guide RNA sequence 5’ GAGGTCATCTCGAAGCCTC 3’ was cloned into pLentiCRISPRv2 (Addgene # 52961; a gift from Feng Zhang) (Sanjana et al., 2014). Infected cells were enriched by treating cells with 3 µg/ml puromycin before sorting and growing clonal cell lines. Rescue with pTLCV2-mScar-ACK was induced by addition of 250 ng/mL doxycycline (Sigma-Aldrich, #D9891) to the growth media 48 hours prior to plating and to the low serum media 4 hours after plating.

### Transferrin Uptake Assay

WT and *Arpc2* KO JR20 cells were plated on 10 µg/ml FN coated 18-well chamber slides (Ibidi, Cat#: 81816) overnight in DMEM with 10% FBS. Prior to the assay, cells were washed 2X with DPBS and incubated in low serum medium (DMEM + 0.2% FBS) for 30 minutes at 37°C. Subsequently, 2.5 µg/mL transferrin-Alexa 488 (Jackson Immuno, Cat#: 015-540-050) in DMEM + 0.2% FBS was added to the cells. After a 10-or 30-minute incubation at 37°C, cells were fixed by adding paraformaldehyde to the media for a final concentration of 4% and incubating at 37°C for 5 minutes. Cells were washed 3x with DPBS and surface bound transferrin was removed using cold acid buffer (0.5% acetic acid, Fisher, Cat#: A38C-212; 0.5M NaCl, Fisher, Cat#: BP358-10, in DPBS) prior to widefield imaging.

### Drug treatments and washout

Cells were initially plated on 10 µg/ml FN coated glass in DMEM with 10% FBS. After 4 hours, the cells were washed 1x with DPBS and the media was replaced with DMEM with 0.2% FBS containing 150 µM CK666, 5 µM XMD8-87, or DMSO for 40 hours prior to imaging. For CK666 washout experiments, cells were then washed 1x with DPBS and media replaced with DMEM with 10% FBS and imaged every 15 minutes via TIRF microscopy for 24 hours.

### Statistical Analyses

All plots were generated in and statistical analyses performed in Prism (Graphpad) unless otherwise noted. Error bars on graphs represent the mean plus or minus the standard deviation. Significance was determined using unpaired Welch’s t-tests, with P values <0.01 represented as *, <0.005 as **, <0.001 as ***, and <0.0001 as ****.

## Acknowledgements

We acknowledge Hiral Patel, Dr. Bhagawat Subramanian, Dr. Priscila Siesser, Michelle Marchan and Max Hockenberry for valuable insight and feedback. Research reported in this publication was supported by the National Institute of General Medical Sciences under award nos. R35GM130312 (to J.E.B.) and R35GM138183 (to J.R.B.). This research was also supported by the Intramural Research Program of the National Heart Lung and Blood Institute, National Institutes of Health (NIH). The contributions of the NIH authors are considered works of the United States Government. The findings and conclusions presented in this paper are those of the authors and do not necessarily reflect the views of the NIH or the U.S. Department of Health and Human Services.

