## Supplementary figures and images for "Arp2/3-mediated turnover of large clathrin lattices is regulated through the tyrosine kinase ACK"

### Sup Fig 1

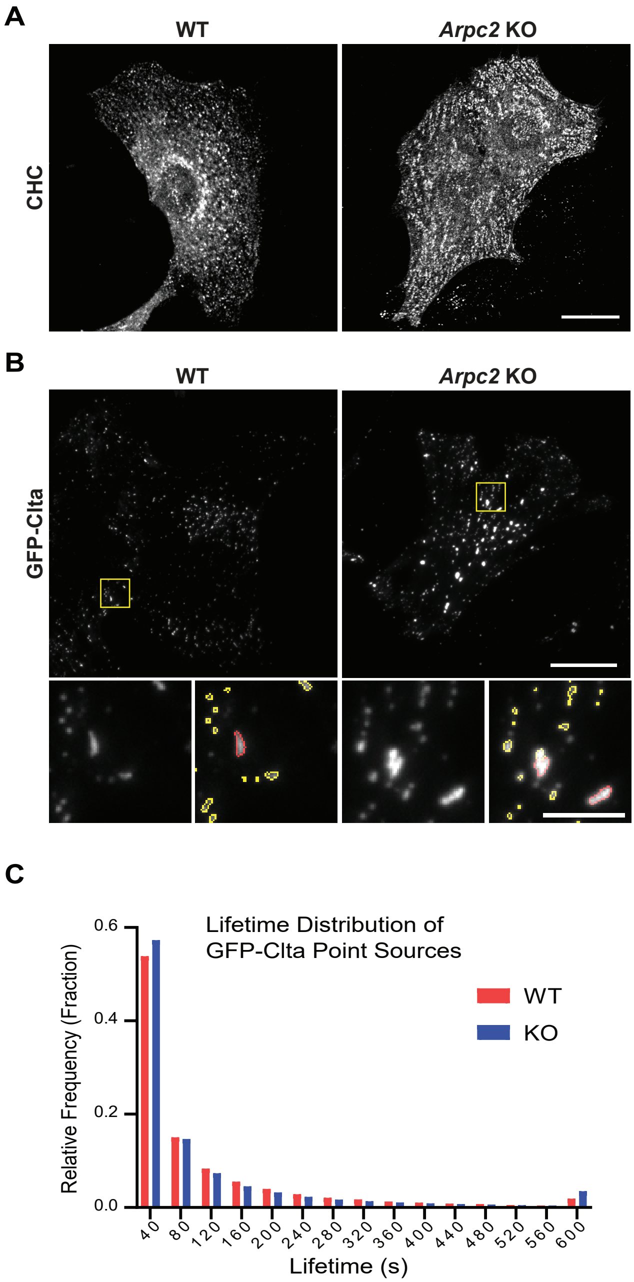

### Sup Fig 2

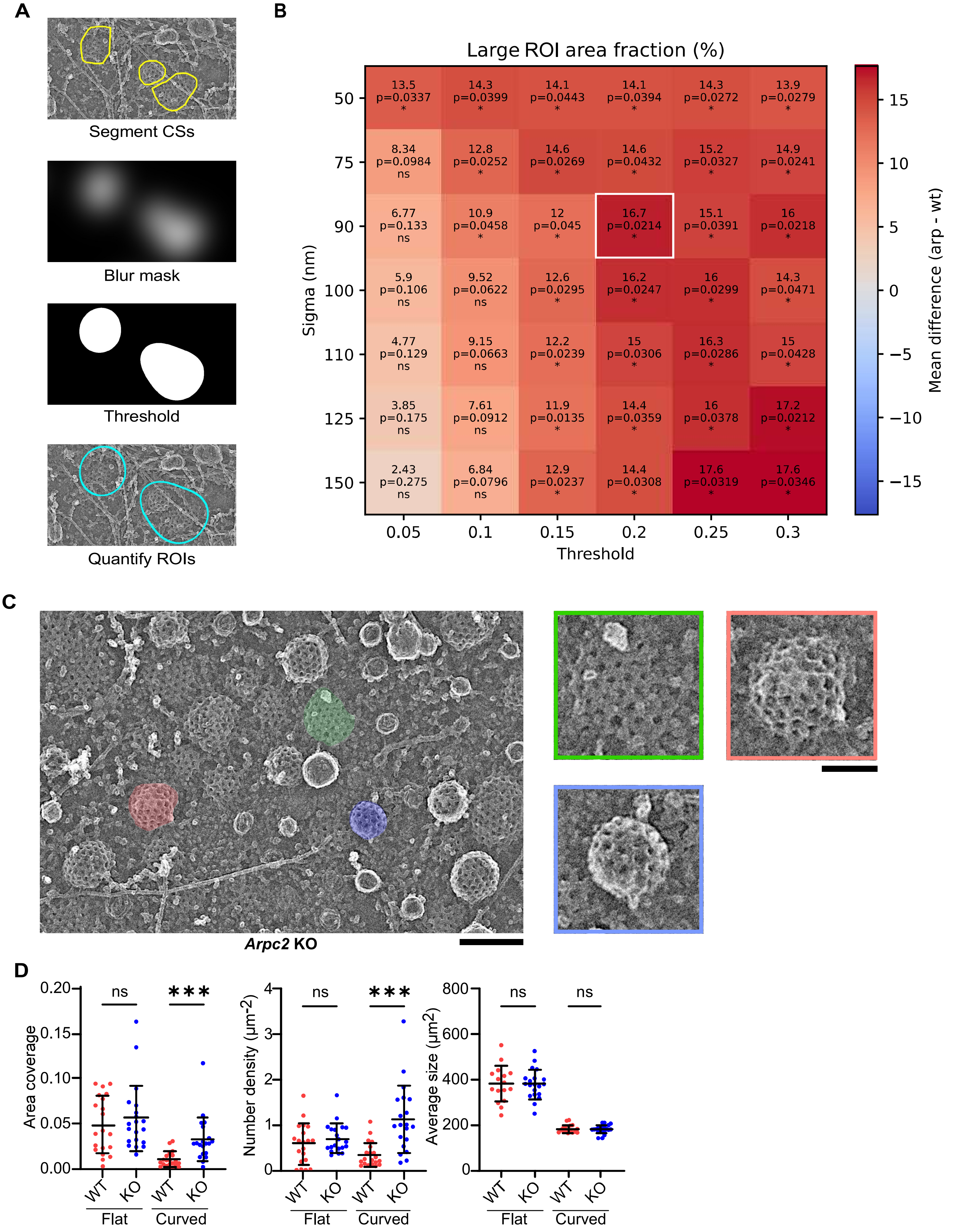

### Sup Fig 3

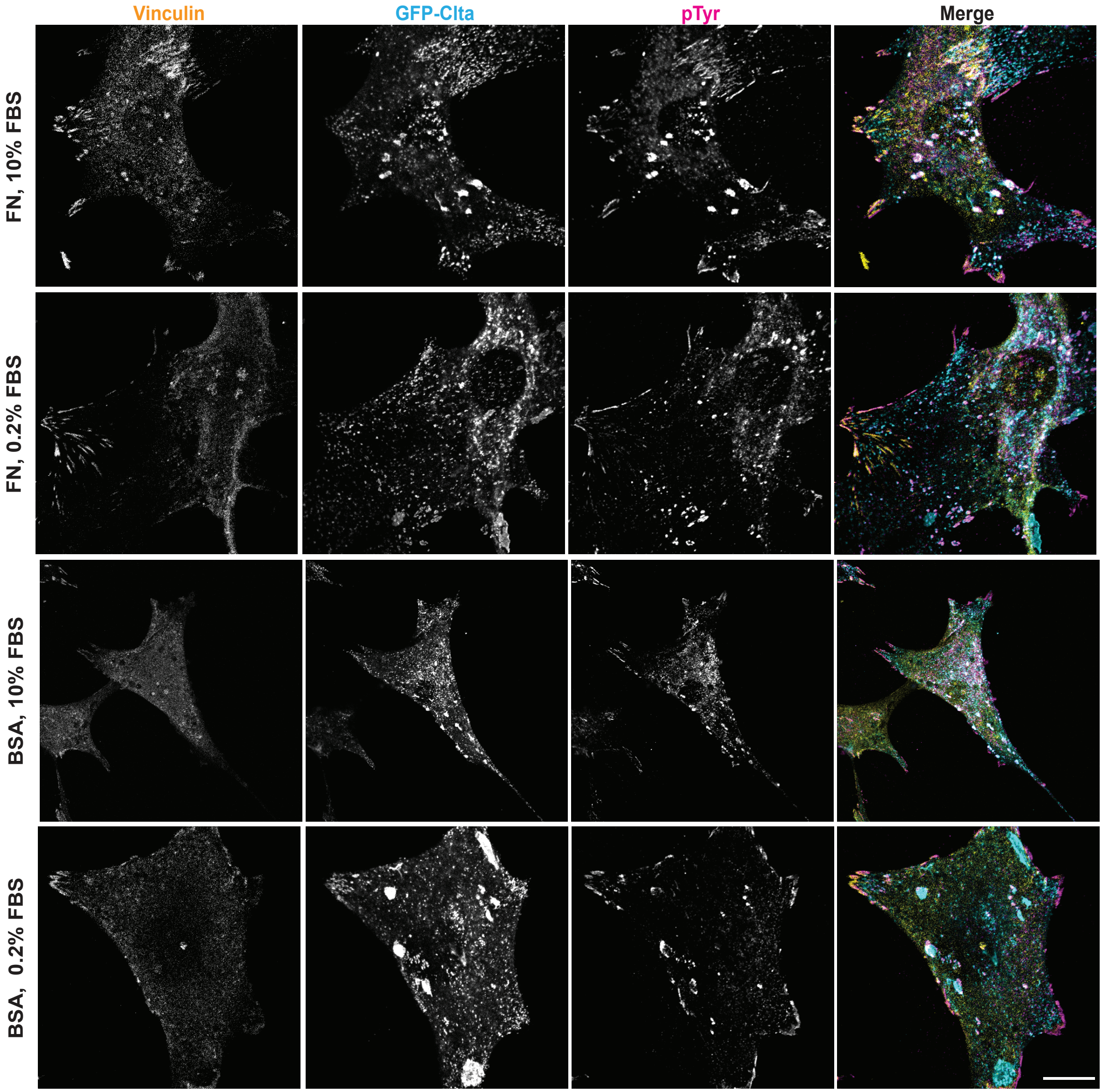
